# Spiral-on-a-Curve: wireless photoacoustic neuromodulation patch

**DOI:** 10.64898/2026.03.04.708963

**Authors:** Shisheng Zhang, Daohuai Jiang, Hengrong Lan, Fei Gao

## Abstract

Focused ultrasound neuromodulation offers a promising noninvasive strategy for precise deep-brain stimulation, yet conventional piezoelectric phased arrays rely on bulky hardware, high-voltage electronics, and complex phase control, limiting their scalability and wearable integration. Photoacoustic approaches enable wireless ultrasound generation but remain constrained by a trade-off between focusing precision, penetration depth, and robustness to optical misalignment. Here, we present a geometrically encoded passive photoacoustic patch (PPP) based on a spherical double logarithmic spiral (SDLS) array that achieves intrinsically stable and programmable acoustic focusing without electronic phase modulation. By distributing hemispherical CNT/PDMS photoacoustic emitters quasi-uniformly over an equal-path spherical surface and orienting each emitter toward a predefined focal point, the device establishes geometry-dominated wavefront convergence. Numerical simulations demonstrate that curved geometry is a prerequisite for phase-free focusing, while the nonperiodic spiral topology suppresses sidelobes and mitigates interference artifacts Compared with continuous spherical or periodic concentric arrays, the SDLS architecture exhibits substantially enhanced robustness to optical axis displacement, reducing focal tilt from > 14° to approximately 5° under 2 mm lateral misalignment. Experimental three-dimensional hydrophone mapping confirms millimeter-scale focusing at approximately 7 mm depth with a full width at half-maximum of 1.3 mm and peak pressures up to 8 MPa under safe laser exposure (≤ 20 mJ/cm^2^). The focal region can be continuously tuned by adjusting illumination aperture size without altering device geometry or excitation schemes. The patch demonstrates excellent thermal and acoustic stability during prolonged operation and enables region-specific motor cortex stimulation *in vivo*, eliciting distinct electromyographic responses in forelimb and hindlimb muscles. By shifting ultrasound beam formation from electronic phase control to intrinsic three-dimensional geometry, this work establishes a lightweight, wire-free, and optically programmable platform for robust wearable neuromodulation and scalable bioacoustic interfaces.

## Introduction

Neuromodulation technologies, which directly intervene in dysfunctional neural circuits, have emerged as transformative therapeutic strategies for neuropsychiatric disorders and metabolic dysregulation, including depression, post-traumatic stress disorder (PTSD), and diabetic neuropathy (*1–3*). Among existing modalities, deep brain stimulation (DBS) offers high spatial precision but is inherently invasive, carrying surgical risks and the potential for irreversible tissue damage (*1, 2*). Noninvasive approaches such as transcranial magnetic stimulation (TMS) and transcranial electrical stimulation (TES) provide improved safety profiles; however, their limited penetration depth and diffuse field distributions restrict effective targeting of deep brain structures, such as the basal ganglia and entorhinal cortex (*4, 5*).

In contrast, ultrasound neuromodulation has gained increasing attention as a next-generation modality capable of noninvasively delivering spatially confined stimulation to deep brain regions with favorable safety characteristics (*4–10*). Despite these advantages, conventional ultrasound systems rely heavily on bulky phased-array hardware and complex multi-channel driving electronics for wavefront shaping. This “heavy-equipment” paradigm not only constrains operation in freely moving conditions but also significantly hinders translation toward lightweight, wearable, or daily-use platforms.

Recent advances in flexible electronics have stimulated the development of wearable ultrasound devices (*11–17*). Pu et al. reported a spiral piezoelectric ultrasound patch capable of beam focusing without complex phase control, demonstrating spatially selective peripheral neuromodulation (*11*). Zou et al. developed a flexible ultrasonic array system for transcranial stimulation and brain therapy, highlighting the benefits of conformal encapsulation and array integration (*17*). Tang et al. introduced a bioadhesive hydrogel-coupled miniaturized ultrasound transducer system that enabled long-term wearable neuromodulation while reducing dependence on conventional coupling media (*12*). Pashaei et al. further integrated multi-channel PZT arrays with on-chip front-end circuits on flexible PCBs to achieve body-conformal imaging–stimulation platforms (*13*).

Although these studies represent important progress in flexible layouts, attachment stability, and functional integration, their fundamental transduction mechanisms remain rooted in thick PZT elements and multi-channel high-voltage electronics. Consequently, power consumption, wiring complexity, and reliance on rigid materials continue to impose substantial limitations on scalability, lightweight deployment, and disposable or large-area wearable implementations. The persistent dependence on rigid piezoelectric materials, high-power electronics, and wired connections fundamentally constrains the realization of low-cost, cable-free, and daily-wear neuromodulation patches (*5*).

To eliminate wired driving architectures entirely, wireless neuromodulation strategies based on the photoacoustic (PA) effect have recently emerged (*18–26*). By converting pulsed optical energy into broadband ultrasound through thermoelastic expansion, photoacoustic systems conceptually enable a fully wireless “light-in, sound-out” paradigm. However, current photoacoustic neuromodulation designs face a critical trade-off between tight focusing and penetration depth or robustness (*18,19,22,23*). For example, Li et al. demonstrated a high–numerical aperture (NA) photoacoustic patch achieving sub–0.1 mm resolution (*19*), yet the design sacrificed penetration depth (limited to hundreds of micrometers) and exhibited pronounced sensitivity to optical illumination conditions. Because acoustic generation in photoacoustic systems is intrinsically coupled to the spatial distribution of optical excitation, small variations in illumination angle, beam uniformity, or lateral alignment can substantially alter the effective emitting aperture, resulting in focal distortion or even loss of focusing.

Therefore, a fundamental gap remains: no wearable photoacoustic neuromodulation platform has yet demonstrated simultaneous millimeter-scale deep focusing, low sidelobe artifacts, robustness to optical misalignment, and programmable focal control without reliance on electronic phase modulation (*4, 5, 18, 19, 27, 28*).

Inspired by the space-filling efficiency of sunflower phyllotaxis patterns (*24,27,28*), we propose a wireless Passive Photoacoustic Patch (PPP) based on a spherical double logarithmic spiral (SDLS) array. In this design, photoacoustic emitters are quasi-uniformly and nonperiodically distributed over an equal-path spherical surface whose center coincides with a predefined focal point. This geometry intrinsically enforces time-of-flight convergence, enabling phase-free geometric focusing. The nonperiodic spiral topology suppresses coherent sidelobe enhancement commonly observed in periodic arrays, thereby improving focal stability and reproducibility within a finite aperture.

Distinct from conventional point-like or periodically sampled photoacoustic emitters, the PPP arranges hemispherical, directionally oriented emitters on a spherical surface centered at the target focus. Each emitter’s axis is aligned toward the focal point, enabling geometric modulation of effective radiation direction while preserving equal-path propagation conditions. In this way, wavefront formation is governed by three-dimensional structural geometry rather than electronic phase delays.

We experimentally demonstrate spatially reproducible millimeter-scale acoustic focusing, supported by comprehensive three-dimensional field mapping and focal characterization. Systematic measurements further reveal that focal position and main-lobe morphology remain stable over a wide range of incident angles and lateral optical axis displacements, indicating strong robustness against illumination misalignment. Moreover, by adjusting the illuminated area of the incident laser beam, we achieve programmable control of focal size and energy distribution without altering device geometry or excitation schemes.

Together, these results establish a structurally simple yet optically programmable platform for generating robust focused ultrasound fields, offering a lightweight and intrinsically stable solution for wearable photoacoustic neuromodulation and next-generation bioacoustic interfaces.

## Results

### Design of the spherical logarithmic-spiral photoacoustic patch

Figure 1A illustrates the envisioned use scenario and operating principle of the passive photoacoustic patch (PPP). The patch conformally interfaces with the skin above a target neural region. Upon irradiation by nanosecond laser pulses, the embedded photoacoustic emitters rapidly absorb optical energy and undergo transient thermoelastic expansion, generating broadband ultrasound. The pressure waves emitted from spatially distributed elements superpose coherently after transmission through the skull and converge at a predefined intracranial target, enabling noninvasive, spatially confined focused-ultrasound neuromodulation. The acoustic transmission of the mouse skull (≈ 69%) provides a physical basis for efficient energy delivery to the brain.

**Figure 1:**
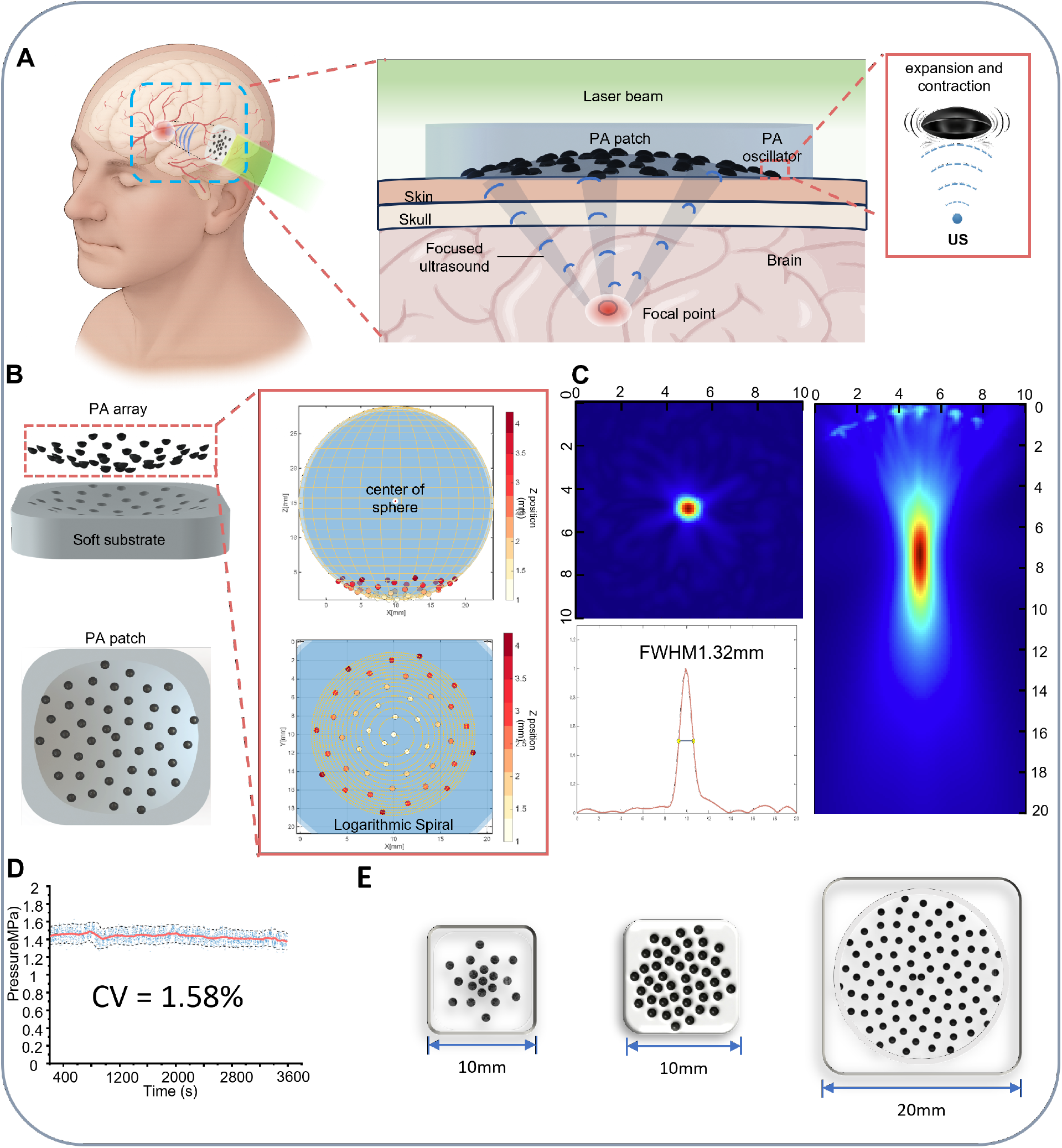
Overview and design of the spherical logarithmic-spiral passive photoacoustic patch (PPP) for neuromodulation. (**A**) Schematic of a fully integrated, stretchable photoacoustic patch and its operating principle under nanosecond pulsed laser excitation for focused-ultrasound stimulation. (**B**) Layout of the PPP: hemispherical photoacoustic emitters embedded in a compliant substrate and arranged along a spherical double logarithmic spiral. (**C**) Simulated acoustic field of the PPP, including focal-plane pressure map and lateral line profile (millimeter-scale full width at half maximum, FWHM), and XZ-plane distribution showing an axial focal depth of approximately 7 mm. (**D**) Long-term output stability under continuous pulsed excitation for 1 h (coefficient of variation, CV = 1.58%). (**E**) Examples of customizable PPP designs with different sizes and emitter counts enabled by scalable manufacturing.

The PPP is built on a transparent polydimethylsiloxane (PDMS) substrate, whose high optical transmittance allows efficient light delivery to the photoacoustic elements. By tuning the base-to-curing-agent ratio, the PDMS modulus can be adjusted to the same order as that of skin, while retaining excellent biocompatibility for wearable applications (Supplementary Information). Each photoacoustic element consists of a carbon nanotube (CNT)/PDMS composite: CNTs efficiently absorb nanosecond optical pulses and convert them into heat, which is transferred to PDMS to drive rapid expansion during pulse-on and fast contraction during pulse-off, facilitated by the high thermal conductivity of CNTs. Because both the emitters and the substrate contain PDMS, covalent crosslinking of siloxane chains during curing enables a monolithic encapsulation, improving mechanical integrity, compliance, and long-term durability—critical for stable operation in physiological environments.

Figure 1B shows the array layout and geometric configuration. We define the basic unit that converts optical energy to ultrasound as a photoacoustic emitter. All emitters are identical hemi-spherical shells (radius 0.3 mm) embedded in an elastomeric substrate, forming a continuous, conformal three-dimensional array. Instead of periodic sampling, the PPP employs a spherical double logarithmic-spiral arrangement that achieves quasi-uniform filling within a finite aperture. Specifically, two intertwined spiral trajectories are constructed in the top view (XY plane) using golden-angle increments (≈ 2.399 rad) with a fixed angular phase offset, producing an interleaved “double-spiral” pattern. The radial coordinate increases with the square root of the element index to maintain approximately uniform areal density. The 2D distribution is then mapped onto a spherical surface in 3D such that all emitter centers lie on an equal-path sphere defined with respect to the prescribed focal point.This construction enforces identical propagation distances from each emitter to the focus, enabling time-of-flight convergence and coherent summation without electronic phase control. In addition, each hemispherical element is directionally oriented toward the target focus to further strengthen geometric focusing. Detailed geometric construction is provided in Methods.

Importantly, the spherical double-spiral architecture is not merely a layout choice but a geometry-optimization strategy inspired by phyllotaxis. It confers multiple performance advantages. (i) Equal-path spherical placement preserves temporal convergence over clinically relevant propagation distances, supporting millimeter-scale lateral resolution and effective focal depths on the millimeter scale or beyond. (ii) By breaking long-range spatial periodicity and symmetry—unlike concentric rings or periodic lattices—the interleaved, aperiodic sampling suppresses coherent sidelobe reinforcement and reduces structured interference artifacts (*27–29*). (iii) The radially graded distribution (denser near the center, sparser toward the periphery) maintains a robust set of contributing emitters under illumination-angle variation or lateral beam misalignment, improving tolerance to optical mismatch. (iv) Because the effective excitation aperture is defined optically, the number of participating emitters can be continuously tuned by adjusting the illuminated area, allowing focal-size modulation without modifying array geometry.

Figure 1C summarizes the simulated acoustic field of the PPP. The focal-plane pressure map shows strong lateral confinement, with a lateral full width at half maximum (FWHM) on the order of approximately 1.3 mm, indicating millimeter-scale focusing. The XZ-plane distribution further shows progressive axial convergence and a stable focal region at approximately 7 mm depth, validating the geometric-focusing capability of the spherical array. To assess long-term stability, we continuously excited the PPP for 1 h using a 1064 nm laser (pulse fluence 20 mJ cm^−2^, pulse width 8 ns, repetition rate 10 Hz). The focal peak pressure exhibited minimal fluctuation with a coefficient of variation (CV) of 1.58% (Fig. 1D), confirming stable acoustic output under prolonged pulsed operation. Owing to low-cost and scalable manufacturing, the PPP can be readily customized in array size, emitter count, and layout to accommodate different targeting requirements (Fig. 1E).

### Geometric rationale and focusing robustness of the spherical spiral PPP

To systematically evaluate how array geometry governs focus formation, stability, and field quality, we performed stepwise comparative simulations in k-Wave under identical medium parameters and photoacoustic excitation conditions (Fig. 2). Because photoacoustic excitation produces broadband pulses for which stable coherent addition via electronic phase control is impractical, we focused on geometric determinants of focusing performance and robustness.

**Figure 2:**
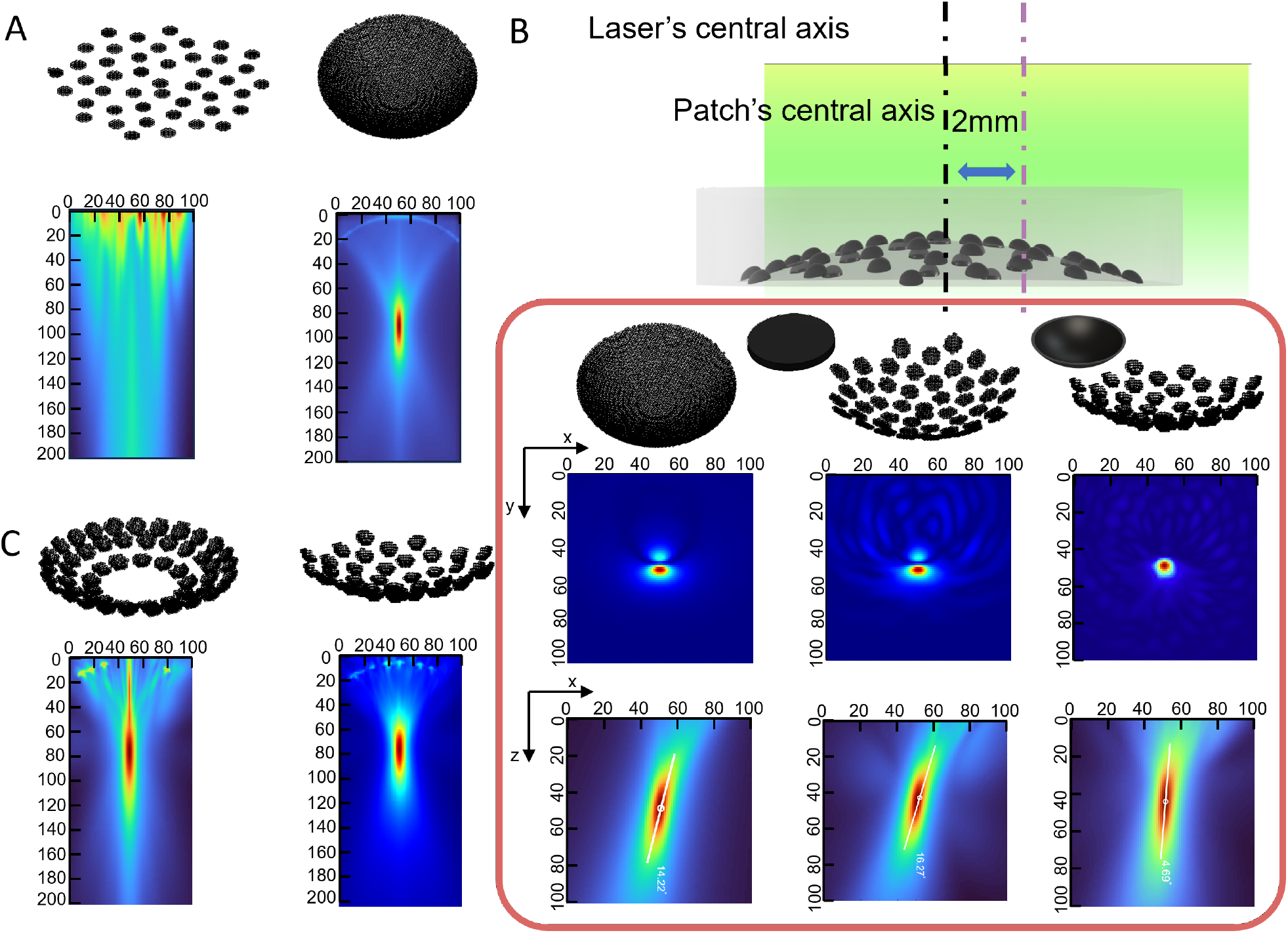
k-Wave simulations of geometric focusing and robustness enabled by the spherical logarithmic-spiral PPP. (**A**) Comparison of a planar logarithmic-spiral disk emitter (left) and a continuous spherical emitter (right) under axial illumination: the planar configuration yields spatially distributed interference without a clear focus, whereas the spherical geometry forms a focal region due to equal time-of-flight to the sphere center. (**B**) Robustness under illumination misalignment: a 2 mm lateral offset is introduced between beam axis and patch center. Three geometries constrained by the same spherical radius are compared—continuous spherical emitter, spherical arc array, and spiral-sampled hemispherical bowl array. XY focal-plane and XZ distributions show pronounced focal distortion and axis rotation for the continuous sphere (14.22°) and arc array (16.27°), whereas the spiral bowl maintains a single focus with modest tilt (5.14°). (**C**) Comparison between a concentric-ring bowl array and a spherical logarithmic-spiral bowl array: RMS pressure maps show structured interference fringes and near-field hotspots for concentric rings, while the spiral array yields a more continuous focal region with reduced off-target structure.

### Curved geometry is required for phase-free photoacoustic focusing

We first compared planar and curved emitter arrangements under ideal illumination alignment (Fig. 2A). When emitters were placed on a plane—even with logarithmic-spiral sampling—the resulting field exhibited discrete interference patterns without a well-defined focus. In contrast, a continuous spherical emitter surface produced a localized high-energy region near the sphere center, consistent with geometric focusing. This difference arises from the equal-path property of spherical geometry: points on the spherical surface share identical propagation distance to the center, enabling synchronous arrival without phase modulation. These results establish curved geometry as a prerequisite for phase-free photoacoustic focusing.

### Emitter discretization and directionality determine robustness under optical misalignment

Having established that curved geometry supports focusing, we next compared the robustness of different curved geometries under lateral beam-axis misalignment (2 mm offset) using a flat-top beam whose area just covered the patch, while keeping the same spherical radius constraint (5 mm) (Fig. 2B). For a continuous spherical emitter, beam offset caused pronounced focal deformation in the XY plane and a clear rotation of the focal axis in the XZ plane (≈ 14.22°), indicating high sensitivity to optical alignment. A spherical-arc array (geometrically matched to the later bowl array, but replacing discrete directional elements with a continuous arc surface) showed similarly severe distortion, with an even larger focal-axis rotation (≈ 16.27°). In contrast, the hemispherical bowl-emitter array sampled by the spherical logarithmic spiral preserved a single, compact focus under the same misalignment, with only modest axis tilt (≈ 5.14°). These comparisons indicate that, given a focus-capable curved geometry, the discretization strategy and directional radiation characteristics critically determine stability to optical mismatch.

### Aperiodic sampling suppresses sidelobes and structured interference

Beyond stability, we assessed off-target energy distribution for different discretization schemes (Fig. 2C). A concentric-ring bowl array produced an axially elongated high-energy region accompanied by strong periodic interference fringes and near-field hotspots, indicating that sidelobe energy tends to concentrate into structured artifacts. The spherical logarithmic-spiral bowl array instead generated a more continuous focal region with smoother boundaries, markedly reduced structured sidelobes, and lower-amplitude, less organized background energy. This supports the conclusion that logarithmic-spiral aperiodic sampling suppresses periodic-array sidelobes and improves overall field quality.

### Experimental characterization and optical programmability of the focal region

After validating the geometry design in simulation, we experimentally mapped the PPP acoustic output using a three-dimensional hydrophone scanning system (Fig. 3A). The system comprised a motorized 3D translation stage, a hydrophone mounted on the stage, an amplifier, an oscilloscope, and a host computer for acquisition and field reconstruction. Time-domain signals were recorded on a predefined spatial grid and processed to reconstruct pressure-amplitude maps on the focal plane and propagation planes.

**Figure 3:**
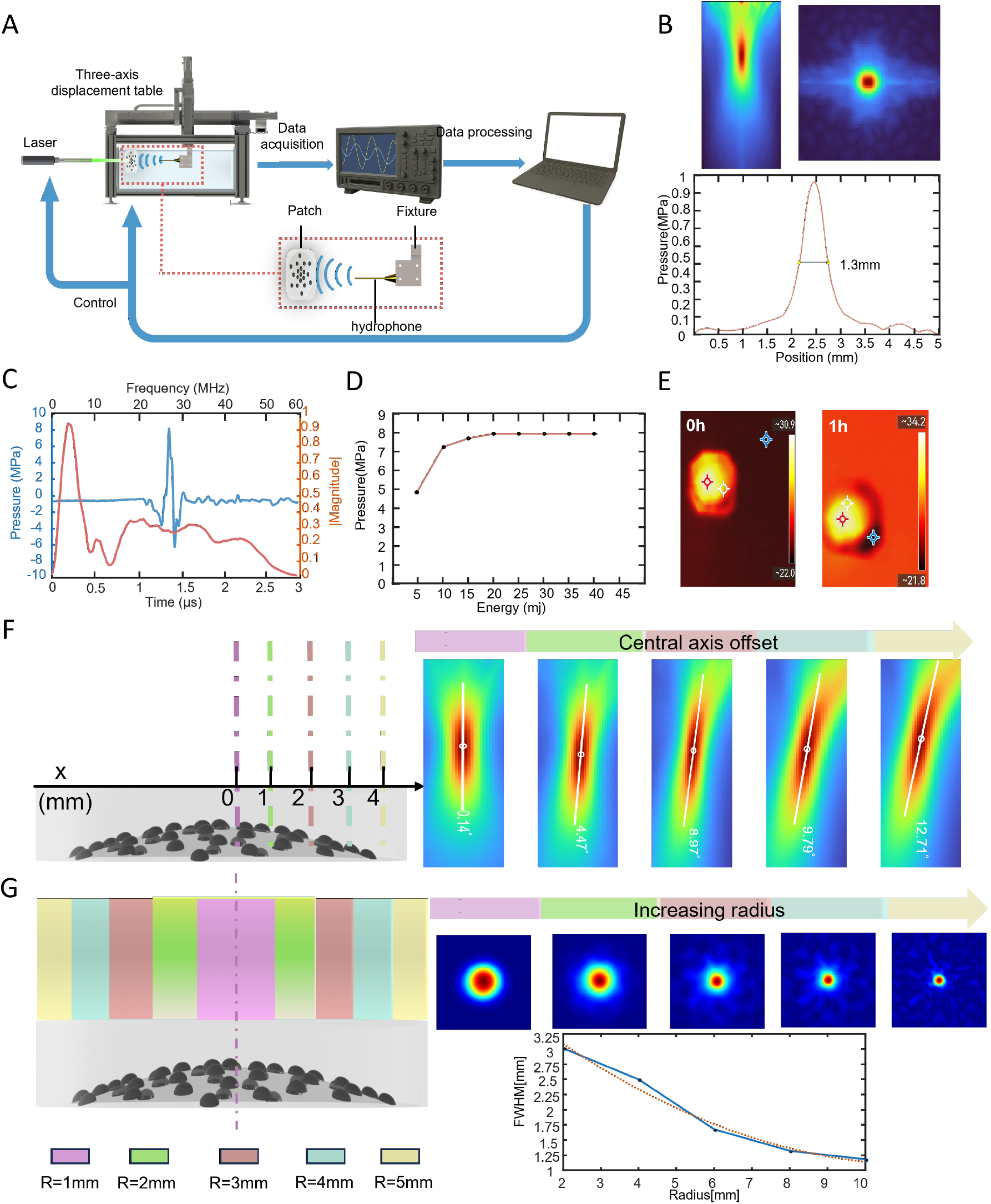
Experimental acoustic characterization and optical programmability of the PPP focus. (**A**) Hydrophone-based 3D scanning system for field acquisition and reconstruction (motorized 3D stage, hydrophone, amplifier, oscilloscope, and computer). (**B**) Measured pressure distributions in the XY focal plane and XZ propagation plane under normal (90°) incidence, yielding a focal-plane FWHM of 1.3 mm. (**C**) Representative time-domain waveform and spectrum at the focus, showing broadband content (0–60 MHz) with a dominant component around 4–5 MHz and peak pressure up. (**D**) Peak pressure versus laser fluence, showing saturation near 20 mJ cm^−2^. (**E**) Infrared thermography before operation (0 h) and after 1 h continuous operation, indicating temperatures within a safe range. (**F**) XY focal-plane pressure maps under increasing lateral beam-axis offsets, showing progressive focal-axis rotation and asymmetric deformation. (**G**) Optical control of focal size by varying beam radius: focal-plane maps and an FWHM–radius trend showing progressive focal contraction with increasing illumination aperture.

Under normal (90°) laser incidence, the PPP produced a clear localized focal region in both the XY focal plane and XZ propagation plane (Fig. 3B), with a focal-plane FWHM of 1.3 mm, confirming millimeter-scale focusing experimentally. Representative time- and frequency-domain analysis at the focus (Fig. 3C) showed a peak pressure up to 8 MPa, a dominant frequency component near 4.05 MHz, and broadband spectral content spanning 0–60 MHz.

To relate acoustic output to optical exposure and assess practical safety margins, we measured peak pressure versus laser fluence (Fig. 3D). At 5 mJ cm^−2^, the peak pressure reached 4.8 MPa. The output increased with fluence and exhibited saturation near 20 mJ cm^−2^, which is within the reported skin safety threshold for the employed wavelength and pulse conditions. We further evaluated thermal safety by operating the patch for 1 h at 40 mJ cm^−2^ (twice the saturation fluence), with 10 Hz repetition, 1064 nm wavelength, and 8 ns pulse width. With coupling gel applied to emulate use conditions, the maximum surface temperature rose from 30.9 °C to 34.2 °C after 1 h, remaining within a safe temperature range. To quantify tolerance to lateral beam-axis deviation commonly encountered in wearable settings, we introduced progressive beam offsets along one lateral direction and reconstructed YZ-plane fields to evaluate focus tilting. The focal axis tilt increased systematically with offset, from near-zero (0.14°) at nominal alignment to 12.71° at the largest tested offset, demonstrating controlled degradation rather than catastrophic focus loss across the misalignment range.

Finally, we demonstrated optical programmability of the focal region by varying the illuminated area. As the beam radius increased over a broad range, the focal spot progressively contracted (Fig. 3G), and quantitative analysis showed a monotonic dependence of FWHM on beam radius.

These results establish that the PPP enables programmable control of focal size and energy distribution solely by adjusting illumination, without modifying device geometry or introducing phased excitation.

### *In vivo* region-specific motor cortex stimulation and EMG validation

To validate region-specific neuromodulation *in vivo*, we stimulated the mouse motor cortex and recorded downstream electromyography (EMG) responses from corresponding limbs (Fig. 4). As shown in Fig. 4A, the mouse was fixed in a stereotaxic frame, and the PPP was integrated with an optical fiber via a rigid holder. The fiber output was placed in direct contact with the patch surface to ensure stable photoacoustic excitation. The patch assembly was mounted on a 3D translation stage, enabling precise positioning on the skull for targeting distinct motor cortical subregions.Stimulation of the forelimb motor cortex elicited a clear time-locked transient EMG response in the forelimb (Fig. 4B), characterized by a distinct spike with short duration, indicating effective activation of the forelimb motor pathway. Stimulation of the hindlimb motor cortex produced a higher-amplitude EMG response with more complex temporal structure (Fig. 4C), including multiphasic components and longer activation duration. The systematic differences between forelimb and hindlimb EMG signatures are consistent with physiological distinctions in muscle volume, motor-unit recruitment, and corticospinal organization, supporting that the recorded signals arise from region-specific neural activation rather than nonspecific mechanical or electrical artifacts.

**Figure 4:**
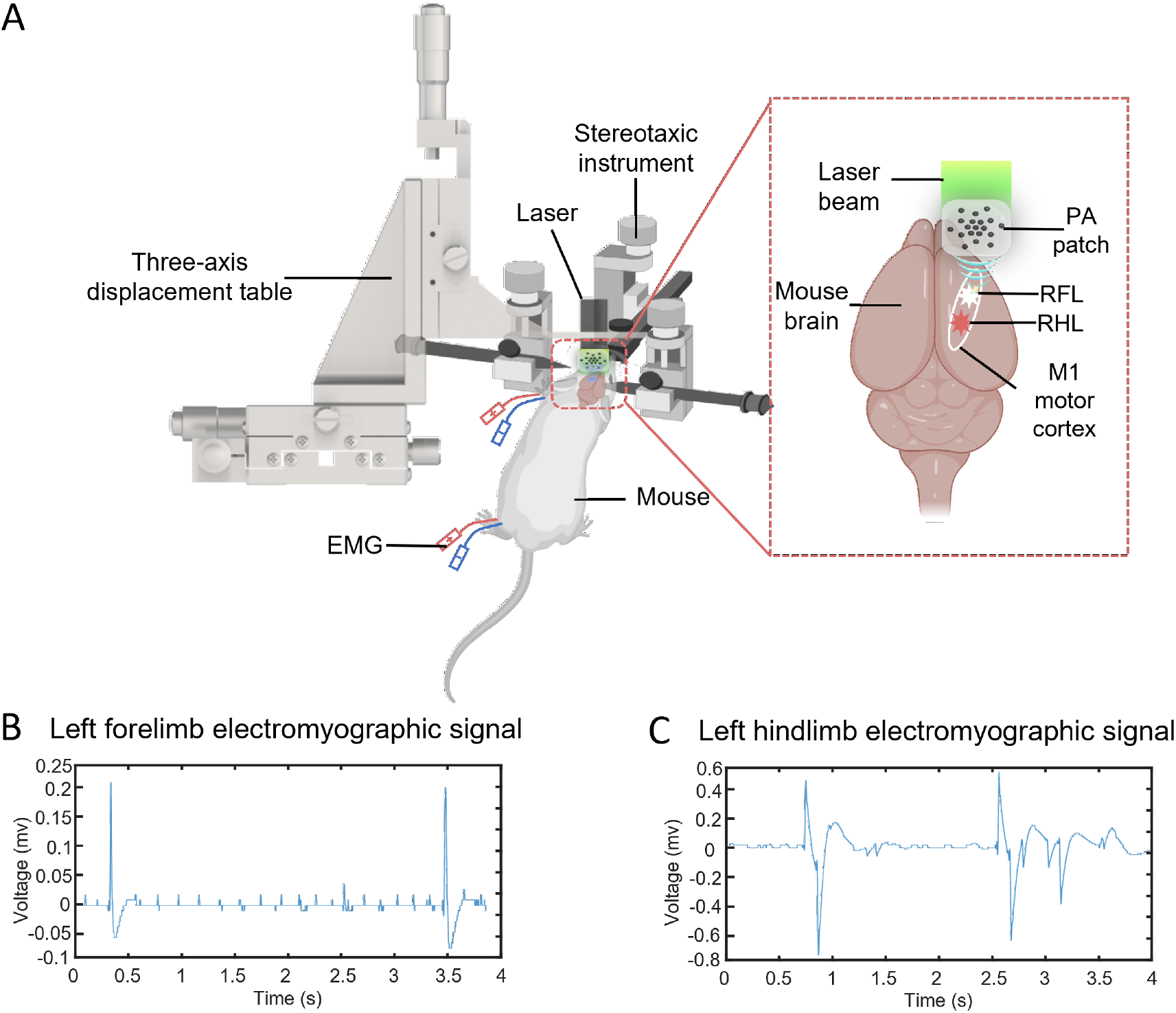
*in vivo* region-specific motor cortex stimulation enabled by the PPP and corresponding EMG responses. (**A**) Experimental setup: mouse fixed in a stereotaxic frame; PPP and optical fiber integrated via a rigid holder; fiber output in contact with the patch for stable excitation; 3D translation stage enables precise positioning over different motor cortical regions. (**B**) Forelimb EMG recording during stimulation of the forelimb motor cortex, showing time-locked transient activation. (**C**) Hindlimb EMG recording during stimulation of the hindlimb motor cortex, exhibiting higher amplitude and more complex waveform structure consistent with differential motor-unit recruitment.

Collectively, these *in vivo* results demonstrate that the spherical logarithmic-spiral PPP can achieve region-specific cortical stimulation under physiological conditions, providing direct biological evidence for its potential in noninvasive neuromodulation.

## Discussion

In this study, we present a geometry-encoded photoacoustic patch based on a CNT/PDMS composite that enables stable and optically programmable acoustic focusing. Unlike conventional phased-array systems that rely on electronically imposed phase delays (*4, 5, 9, 13*), our approach exploits a spherical double logarithmic spiral topology to directly regulate the spatiotemporal distribution of photoacoustic excitation at the device-structure level. In this configuration, the acoustic wavefront is intrinsically shaped by three-dimensional geometry, shifting the mechanism of ultrasound focusing from externally controlled phase modulation to internally defined geometric convergence.

A defining feature of the proposed platform is that acoustic focusing is determined entirely by spatial geometry rather than synchronized electrical excitation. As a result, stable millimeter-scale focal regions can be achieved without multi-channel driving electronics, phase calibration, or complex wiring. Both numerical simulations and experimental field mapping confirm that the structure generates reproducible three-dimensional focal spots, validating the feasibility of geometry-dominated focusing in broadband photoacoustic systems.

An important consequence of this strategy is its intrinsic tolerance to illumination variability. Systematic evaluation of incidence angles (0–75°) and lateral beam-axis displacements (0–4 mm) demonstrates that, within the experimentally tested range, focal position and main-lobe morphology remain preserved without focus splitting or pronounced sidelobe amplification. In contrast to electronically phased arrays, where minor phase errors can severely degrade focusing performance, optical excitation in the present design functions solely as an amplitude weighting of a geometrically predefined acoustic wavefront (*24, 25*). Variations in illumination therefore modulate the effective excitation aperture but do not disrupt the equal-path condition that governs temporal convergence. This geometry-dominated, optically weighted mechanism enables stable focusing under moderate misalignment conditions.

Beyond robustness, the platform enables optical programmability of the focal domain. Although focal position and phase coherence are fixed by array geometry, the effective aperture can be continuously adjusted through illumination area modulation, allowing dynamic control over focal size and energy distribution without altering device architecture or excitation circuitry. This decoupling of structural complexity from acoustic controllability provides a compact and efficient pathway for adaptive ultrasound manipulation.

Compared with traditional piezoelectric arrays (*11–13, 15–17*), the proposed patch eliminates large-scale electrode networks and high-voltage driving electronics, substantially reducing system complexity. The absence of metallic electrodes also offers potential compatibility with electro-magnetically sensitive environments such as magnetic resonance imaging (MRI). Furthermore, the use of soft materials and mold-based fabrication supports scalability and customizable geometries across different anatomical targets.

It is worth noting that large numerical apertures, while enabling tighter focusing, may introduce wavefront distortion in acoustically heterogeneous media. However, because the acoustic phase profile of this platform is fully defined by surface geometry, such distortions could be addressed through geometry-level precompensation. Integration with individualized imaging and inverse-design frameworks may enable subject-specific structural optimization to mitigate propagation-induced aberrations.

In summary, this work establishes a geometry-encoded photoacoustic ultrasound platform that integrates spatial precision, intrinsic robustness, and optical programmability within a lightweight architecture. By decoupling wavefront formation from electronic phase control, the proposed approach provides a new design paradigm for compact, scalable, and multifunctional ultrasound systems, with potential applications in precision neuromodulation, acoustic manipulation, and broader biomedical interfaces.

## Methods

### Geometry design of the spherical double logarithmic spiral PPP

The passive photoacoustic patch (PPP) consists of 100 identical hemispherical-shell photoacoustic emitters embedded within a transparent PDMS substrate. The centers of all emitters are constrained to lie on a common spherical surface centered at a predefined focal point

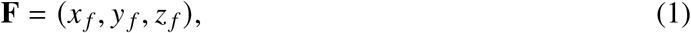

defined in a three-dimensional Cartesian coordinate system whose origin is located at the geometric center of the patch base. Here, *x*_*f*_ and *y*_*f*_ denote the lateral focal coordinates within the patch plane, and *z*_*f*_ specifies the axial focal depth along the surface normal direction. By enforcing this equal-path spherical constraint, geometric time-of-flight convergence at **F** is achieved without electronic phase modulation.

### Planar sampling using an interleaved golden-angle logarithmic spiral (XY)

To achieve quasi-uniform filling within a finite aperture while suppressing sidelobes and interference artifacts associated with periodic sampling, a bidirectionally interleaved “sunflower-like” (phyllotaxis) logarithmic spiral distribution is adopted in the top-view (XY) plane.

The total number of emitters is defined as

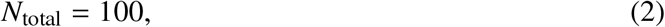

which are generated from two interleaved logarithmic spirals, each containing

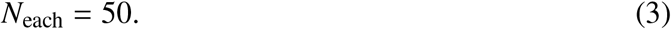

For clarity, we introduce a radial index *k* = 0, 1, …, *N*_each_ − 1 and a spiral index *m* ∈ {1, 2}.

### Radial sampling

The common radial coordinate at index *k* is defined as

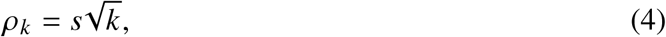

where *s* is a scaling factor controlling the maximum array radius. This formulation ensures that at each radial level *k*, two emitters—one from each spiral—are placed at the same radius *ρ*_*k*_, thereby preserving radial consistency while allowing angular interleaving.

### Angular sampling (interleaved spirals)

The golden angle is defined as

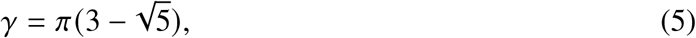

which is known to minimize angular periodicity and promote uniform spatial filling (*30*).

At radial index *k*, the polar angles of the two spirals are defined as

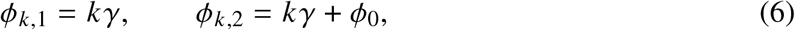

where *ϕ*_0_ = *γ*/2 introduces a fixed angular offset between the two spirals. This offset interleaves the angular positions of the two sequences, further disrupting residual symmetry associated with a single spiral and reducing long-range periodic correlations.

### Planar coordinate representation

Using the substrate center (*x*_*c*_, *y*_*c*_) as the planar origin, the coordinates of the sampling points are expressed as

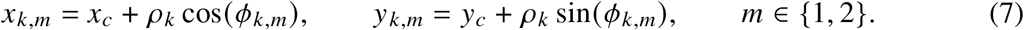

The points generated from both spirals are then merged in ascending radial order to form the final set of 100 emitter positions in the XY plane, constituting a bidirectionally interleaved golden-angle logarithmic spiral distribution.

### Mapping from planar sampling to a spherical (bowl-shaped) array (XYZ)

To realize geometric focusing with a curved aperture, the planar sampling points generated in the XY plane are mapped onto a common spherical surface such that all emitter centers satisfy an equal-distance constraint with respect to the predefined focal point. Let the sphere radius be *R*. For each planar coordinate (*x*_*i*_, *y*_*i*_), the corresponding three-dimensional emitter center (*x*_*i*_, *y*_*i*_, *z*_*i*_) is computed as

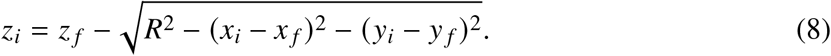

Here, the branch is selected such that the array lies on the outer (upper) hemisphere relative to the focus.

If the expression under the square root becomes negative (i.e., the planar coordinate falls outside the valid spherical projection domain), the value is clipped to zero to ensure geometric feasibility. Through this mapping, all emitter centers **P**_*i*_ = (*x*_*i*_, *y*_*i*_, *z*_*i*_) lie on the same spherical surface centered at **F**, forming a bowl-shaped aperture surrounding the focal point.

### Orientation and geometric parameterization of hemispherical-shell emitters

To ensure that the radiation direction of each photoacoustic emitter aligns with the geometric focusing direction, all hemispherical-shell emitters are designed with their openings facing the predefined focal point **F**.

For the *i*-th emitter, the center position is defined as

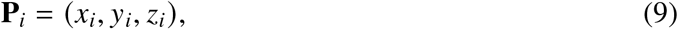

where **P**_*i*_ represents both the geometric center of the hemispherical shell and its spatial placement within the array. As all **P**_*i*_ lie on a sphere centered at **F**, the equal-path propagation condition is inherently satisfied.

The unit direction vector pointing from the focus to the emitter center is defined as

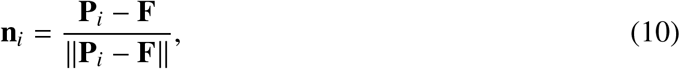

which specifies the axial normal direction of the hemispherical-shell emitter.

The inner surface of each hemispherical shell is formed by a spherical segment of radius *r*_*b*_ centered at **P**_*i*_, with its aperture normal aligned with **n**_*i*_. This ensures that the primary radiation direction of every emitter is precisely oriented toward the predefined focal point **F**. The aperture diameter of each hemispherical shell is 2*r*_*b*_.

Through this geometric construction, all emitters share identical curvature radius and geometric dimensions while collectively forming a spherical photoacoustic array with intrinsic geometric focusing capability.

### Numerical simulations

Three-dimensional acoustic wave propagation simulations were performed using the `k-Wave` tool-box in MATLAB (*31*). A Cartesian voxel grid was employed with perfectly matched layer (PML) absorbing boundaries of thickness 10 grid points (`PMLInside = false`). The grid dimensions were *N*_*x*_ = *N*_*y*_ = 100 and *N*_*z*_ = 200, with spatial resolution *d*_*x*_ = *d*_*y*_ = *d*_*z*_ = 1 × 10^−4^ m, corresponding to a computational domain of approximately 10 × 10 × 20 mm^3^.

The propagation medium was modeled as homogeneous water with sound speed *c*_0_ = 1500 m s^−1^ and density *ρ*_0_ = 1000 kg m^−3^. The temporal grid was automatically generated using `kgrid. makeTime(` *c*_0_).

### Photoacoustic source modeling and array–geometry implementation

In the numerical simulations, photoacoustic excitation was modeled using an equivalent initial pressure source formulation. This approach avoids explicitly incorporating optical propagation modeling and ensures that acoustic field formation is primarily governed by the array geometry and the spatial distribution of the incident optical field.

The PPP array consists of *N*_total_ = 100 hemispherical-shell photoacoustic emitters. The bidirectionally interleaved golden-angle logarithmic spiral arrangement in the planar (XY) domain and the spherical geometric construction follow the procedure described in the preceding geometry-design section. Briefly, a dual-spiral “sunflower-like” (phyllotaxis) sampling pattern is first generated in the XY plane. The golden angle is defined as

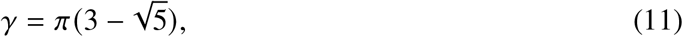

with an angular phase offset between the two spirals given by *ϕ*_0_ = *γ*/2. The radial coordinate follows the scaling law 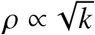, ensuring quasi-uniform filling within a finite aperture.

The resulting two-dimensional sampling points are then mapped onto a spherical surface centered at the predefined focal point

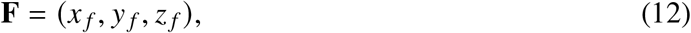

yielding the three-dimensional emitter centers

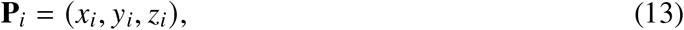

thereby satisfying the equal-path geometric focusing condition. The unit vector pointing from the focal point **F** to each emitter center **P**_*i*_ is defined as

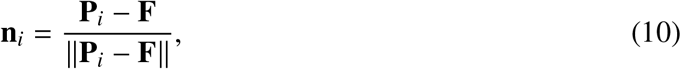

which specifies the axial normal direction of the hemispherical-shell emitter.

In the numerical implementation, the focal position was explicitly set to

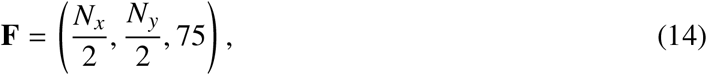

where *N*_*x*_ and *N*_*y*_ denote the lateral grid dimensions of the computational domain, and the third component represents the axial focal depth (in grid units) along the patch normal direction.

Each hemispherical-shell emitter was generated in `k-Wave` using the `makeBowl` function, parameterized by its center **P**_*i*_ and curvature radius *r*_*b*_. The individual bowl masks were combined via logical union to construct the overall three-dimensional source mask, which was subsequently used for acoustic wave propagation simulations.

### Geometric definition of the beam axis

In a three-dimensional Cartesian coordinate system (*x, y, z*), the beam axis is defined to pass through a reference point (*x*_0_, *y*_0_, *z*_0_), where *x*_0_ and *y*_0_ represent the lateral position of the beam center on the reference plane and are used to describe translational offsets relative to the patch center. The axial coordinate *z*_0_ specifies the beam reference height (set to the entrance plane of the computational domain in simulations).

The propagation direction of the beam is parameterized by polar angle *θ* and azimuthal angle *ϕ*. The corresponding unit direction vector is defined as

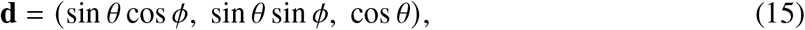

and normalized to satisfy ∥**d**∥ = 1. When *θ* = 0, the beam is normally incident along the patch normal; nonzero *θ* corresponds to tilted illumination.

### Shortest distance from a voxel to the beam axis

For any voxel point **r** = (*x, y, z*) within the computational domain, the displacement vector relative to the beam reference point is defined as

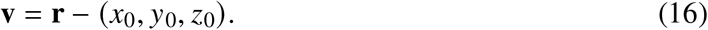

The projection of this vector onto the beam direction **d** is

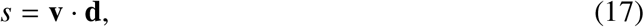

and the perpendicular component is

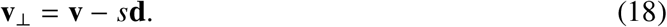

The shortest distance from the voxel to the beam axis is therefore

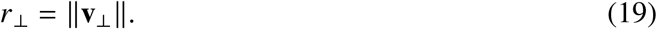

This distance is subsequently used to construct the optical amplitude weighting function.

### Gaussian beam weighting and equivalent initial pressure distribution

Based on the geometric relationship above, the incident optical field is modeled as a Gaussian beam profile (*32*):

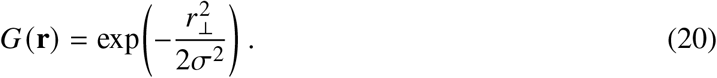

where *σ* denotes the standard deviation controlling the lateral beam width. For consistent comparison across parameter conditions, *G*(**r**) is normalized within the computational domain such that its maximum value equals 1.

The equivalent initial pressure distribution for photoacoustic excitation is then defined as

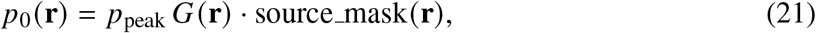

where source mask(**r**) represents the three-dimensional source mask determined by the hemispherical-shell array geometry, and *p*_peak_ is the peak initial pressure amplitude (relative units).

By varying the beam reference point (*x*_0_, *y*_0_), lateral beam-axis displacement within the patch plane can be simulated. Adjusting the directional parameters *θ* and *ϕ* enables modeling of arbitrary tilt angles and azimuthal orientations. This formulation provides a unified description of spatial beam position and direction without explicitly modeling optical propagation, allowing systematic evaluation of focusing robustness under illumination misalignment conditions.

### Sensor plane configuration and acoustic-field reconstruction metrics

To visualize the two-dimensional acoustic field distribution in the focal region, two planar sensor surfaces were defined. The first was a transverse plane at *z* = 75 (XY plane), corresponding to the focal plane for evaluating lateral pressure distribution. The second was a longitudinal plane at *x* = *N*_*y*_/2 (XZ plane), used to examine axial propagation and focal depth structure. The recorded physical quantity was acoustic pressure *p*.

Three-dimensional acoustic wave propagation was solved using the `k-Wave` toolbox, specifically the time-domain solver `kspaceFirstOrder3D`. The resulting time-resolved pressure field *p* (**r**, *t*) was post-processed to derive energy- or RMS-type metrics that more robustly characterize broadband pulsed focusing.

The time-integrated acoustic energy map was computed as

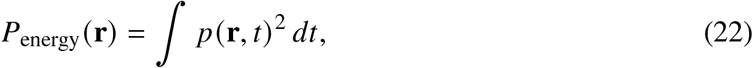

and subsequently normalized. The main-lobe profile was extracted from the normalized energy map, and the lateral focal spot size was quantified using the full width at half maximum (FWHM).

### Quantification of focal tilt using principal component analysis (PCA)

To quantitatively evaluate focus rotation induced by illumination tilt or lateral misalignment, the energy map in the XZ plane was first thresholded (e.g., selecting regions with normalized amplitude greater than 0.5) to isolate the dominant focal region. A weighted covariance matrix was then computed using the energy amplitude as weights. Principal component analysis (PCA) was applied to extract the dominant principal axis vector **v** = [*v*_*x*_, *v*_*z*_]^T^ (*33*).

The tilt angle relative to the *z*-axis was defined as

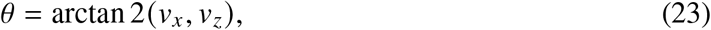

yielding the focal-axis deviation angle. This metric enables quantitative comparison of focal rotation under different geometric configurations or illumination conditions. The extracted principal axis was additionally overlaid on the acoustic-field map to visually validate the numerical estimation.

### Fabrication of the patch substrate and mold

#### 3D model generation and STL export

To fabricate a mold featuring both a spherical-cap outer surface and an array of concave cavities, a three-dimensional geometric model was automatically generated in MATLAB. Boolean subtraction at the point-cloud level followed by *α*-shape surface reconstruction was used to produce an STL file for 3D printing. The procedure is summarized below.

#### Initialization of the base solid

A rectangular base block was first constructed as the initial solid entity, with dimensions *x*_size_ × *y*_size_ × *z*_size_ (e.g., 10 × 20 × 4 mm^3^). The geometric center of the block was set to (*x*_*c*_, *y*_*c*_, *z*_*c*_).

#### Determination of cavity primitive positions

To generate surface cavities corresponding to the emitter array, array point coordinates were first computed in an independent numerical coordinate system. These coordinates were then transformed into the mold-design coordinate system at physical scale. At each transformed location, a small sphere was placed to serve as a negative primitive for cavity generation.

#### Generation of array points and spherical-cap mapping

Within a computational grid defined by *N*_*x*_ = *N*_*y*_ = *N*_*z*_ = 100, planar sampling points (*x*_*i*_, *y*_*i*_) were generated in the XY plane. To obtain the global spherical-cap profile, each planar point was mapped onto a curved surface satisfying the spherical constraint defined by a prescribed focal point **F** and a fixed constraint radius *R* (determined by the distance between a reference point and the focus). The corresponding height coordinate *z*_*i*_ was computed, yielding three-dimensional array points (*x*_*i*_, *y*_*i*_, *z*_*i*_).

This step ensures that all array points lie on the same spherical cap, providing consistent geometric reference positions for subsequent cavity formation in the mold.

#### Coordinate scaling and axial offset

Because the array coordinates were generated in a normalized numerical domain, a coordinate transformation was applied to adapt them to the physical mold scale. A uniform scaling factor (e.g., scale = 0.1) was applied to all three spatial dimensions. Additionally, a constant axial offset was added in the *z*-direction to position the array points within the appropriate depth range of the base block. This guarantees that the subsequent Boolean subtraction produces cavities of appropriate depth on the mold surface.

#### Placement of cavity primitives

After coordinate transformation, a small sphere of radius *r*_*s*_ (e.g., *r*_*s*_ = 0.3) was placed at each array point. These spheres function as negative primitives; during the Boolean subtraction step, they are removed from the base block to form concave cavities corresponding to the emitter positions.

#### Formation of the spherical-cap outer surface

To form the overall spherical-cap (curved) outer profile on top of the cavity array, a larger sphere was introduced as a second subtractive geometric primitive in the mold-design coordinate system. This large sphere defines the global curvature boundary of the mold surface.

The large sphere used to define the global spherical-cap profile is specified by its center and radius:

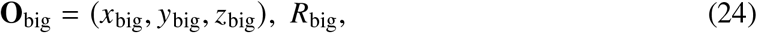

In the implemented example, representative parameters were:

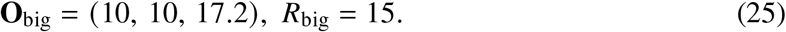

This large sphere defines the outer curved boundary of the mold. All base points located inside the sphere are removed, thereby forming a spherical-cap geometry from the original rectangular block.

#### Point-cloud sampling of the base volume

To perform geometric Boolean subtraction at the point-cloud level, the interior of the rectangular base block is uniformly sampled to generate a point set:

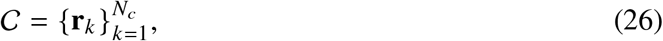

where each point is defined as **r**_*k*_ = (*x*_*k*_, *y*_*k*_, *z*_*k*_). Let the full set of base points be denoted by ℬ.

#### Boolean subtraction: cavity array

For the cavity generation, the *i*-th small sphere (negative primitive) is defined as

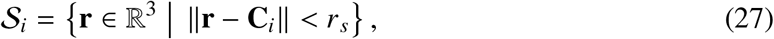

where **C**_*i*_ denotes the center of the *i*-th cavity sphere and *r*_*s*_ is its radius.

The union of all cavity spheres is

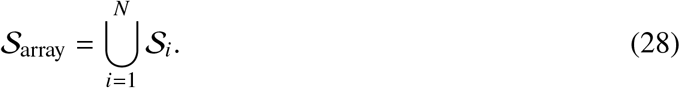

The subtraction of the cavity array from the base block is defined as

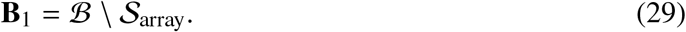

This operation produces an array of concave cavities on the base surface.

#### Boolean subtraction: spherical-cap boundary

The large sphere defining the spherical-cap boundary is described as

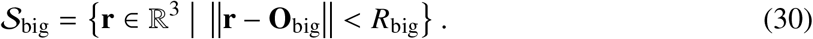

The final retained point set after spherical subtraction is

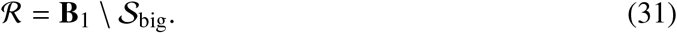

Equivalently, the complete Boolean operation can be expressed as

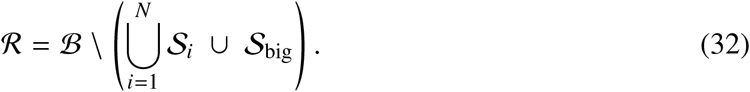

The resulting point cloud ℛ represents the composite geometry consisting of both the spherical-cap outer surface and the cavity array.

#### *α*-shape surface reconstruction and STL export

Because the Boolean operations are performed at the point-cloud level, a continuous surface must be reconstructed prior to STL export.

Let the retained point cloud be

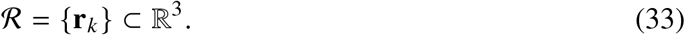

An *α*-shape (*α*-complex) operator 𝒜_*α*_ (ℛ) is applied to reconstruct the outer boundary triangulation. The parameter *α* controls the level of geometric detail (a representative value in our implementation was *α* = 0.2):

- Smaller *α*: preserves fine geometric features
- Larger *α*: produces a smoother outer envelope

The boundary triangular facets are extracted as

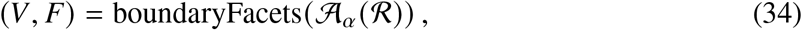

where:

- *V* = {**v**_*i*_ } denotes the vertex set
- *F* denotes the triangular face index matrix

A triangulated surface object is then constructed as

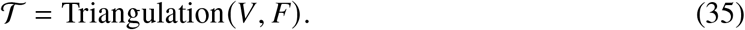

This triangulated mesh 𝒯 represents the reconstructed outer surface of the mold geometry.

#### Laplacian smoothing

To improve mesh quality and reduce potential 3D-printing artifacts while preserving topology, Laplacian smoothing is applied to the triangulated mesh.

Let 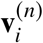 denote the position of the *i*-th vertex at iteration *n*, and let 𝒩 (*i*) denote the set of its neighboring vertices. The iterative update rule is:

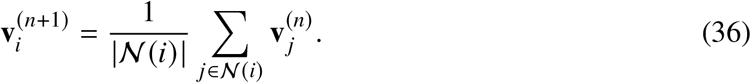

Here, | 𝒩 (*i*) | denotes the number of adjacent vertices. In practice, 5–20 iterations are applied to enhance surface smoothness while maintaining geometric fidelity.

The smoothed triangulation 𝒯 is subsequently exported as an STL file for 3D printing and mold fabrication. This smoothing procedure improves surface smoothness and enhances STL mesh quality while preserving the overall geometric topology.

The smoothed triangulation object was subsequently exported as an STL file:

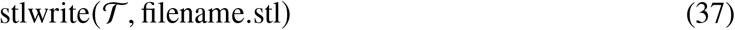

for subsequent 3D printing and mold fabrication.

### Mold fabrication and surface treatment

Based on the reconstructed three-dimensional model, the mold structure was finalized using Bambu Studio and fabricated via 3D printing to obtain a rigid master mold. The printed mold was ultrasonically cleaned in isopropanol (IPA) for 5 min to remove surface contaminants, dried with nitrogen, and placed in an oven for moisture removal prior to casting.

### Substrate fabrication (replication and curing)

To obtain a high-quality spherical-cap substrate with reliable demolding performance, a two-step replication process was employed for PDMS base fabrication.

#### Primary mold fabrication (3D-printed master)

The STL file generated in MATLAB was used to 3D print a rigid master mold matching the target patch geometry. The master mold contained both the global spherical-cap profile and convex features corresponding to the array cavities, serving as the template for subsequent replication.

#### First replication: Ecoflex compliant mold

The 3D-printed master mold was placed in a casting container, and Ecoflex 00-30 elastomer was poured and cured at room temperature or slightly elevated temperature. After curing, the Ecoflex layer was demolded to obtain a flexible positive mold replicating the desired geometry.

The Ecoflex mold offers several advantages:

- High elasticity, facilitating demolding
- Accurate replication of fine surface features
- Reduced stress concentration and tearing during subsequent PDMS casting

This step enables the transition from a rigid printed master to a compliant intermediate mold.

#### Second replication: PDMS substrate fabrication

The Ecoflex mold was then used as the final casting mold. PDMS (Sylgard 184, base:curing agent = 10:1 by mass) was poured onto the mold, degassed under vacuum, and thermally cured (e.g., 60–80 °C for several hours). After curing and demolding, a transparent flexible PDMS substrate was obtained.

The resulting substrate featured:

- The global spherical-cap outer geometry
- An array of concave cavities precisely corresponding to emitter positions

This structure provides accurate spatial alignment and mechanical support for subsequent construction of the photoacoustic emitters within the cavities.

### Material compatibility and interfacial bonding

Because both the substrate and the photoacoustic composite contain PDMS, covalent crosslinking between siloxane chains occurs during secondary casting or local filling. This results in monolithic interfacial bonding, enhancing overall mechanical stability and long-term durability of the device.

### Preparation of CNT/PDMS photoacoustic composite

The CNT/PDMS composite was prepared using an ultrasonication-assisted solution blending method. Multiwalled carbon nanotubes (1.0 g; diameter ∼8 nm; length 10–30 μm) were dispersed in 80.0 g anhydrous ethanol via magnetic stirring (300 rpm, 10 min), followed by probe ultrasonication in an ice bath (300 W, pulsed mode, 15 min).

Subsequently, 10 g of PDMS prepolymer was added (CNT:PDMS mass ratio = 1:10), and stirring and ultrasonication were repeated to improve dispersion uniformity. Ethanol was evaporated in a 70 °C water bath. After solvent removal, PDMS curing agent was added at a 10:1 mass ratio (prepolymer:curing agent), mixed for 10–20 min, and vacuum-degassed for 5–10 min to obtain the CNT/PDMS composite for photoacoustic emitter fabrication.

### Aerosol jet printing and monolithic encapsulation

Because the emitters are concave hemispherical cavities arranged in three-dimensional space, conventional spin coating would lead to edge accumulation and insufficient coverage in deep regions. Therefore, aerosol jet printing (Aerosol Jet 200, Optomec) was employed to conformally deposit the CNT/PDMS composite onto the curved emitter surfaces.

During curing, crosslinking between PDMS chains in the composite and substrate results in monolithic encapsulation, improving mechanical robustness and structural durability.

### Acoustic field generation and three-dimensional scanning characterization

#### Signal acquisition and synchronization

Acoustic pressure signals were acquired in the time domain using a digital oscilloscope. Communication and automated data acquisition were implemented via a USB-VISA interface controlled in MATLAB. The oscilloscope waveform mode was set to Normal, and data were exported in ASCII format.

At each spatial scan position, complete time-domain waveforms from two channels were recorded and stored as independent samples for subsequent analysis. Acquisition was performed in cycles at approximately 0.1 s intervals, corresponding to a sampling frequency of approximately 10 Hz. The total number of acquisitions was monitored in real time to ensure data continuity during scanning.

#### Data compression metric and preprocessing

To extract a scalar metric representing local acoustic field strength at each scan position, feature extraction was performed on the time-domain pressure waveform. The maximum absolute pressure amplitude was used as the intensity metric.

Specifically, for each waveform *p* (*t*), the intensity was defined as

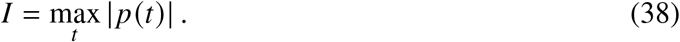

The resulting intensity values were then normalized across the entire scan plane to enable comparison under different illumination or geometric conditions.

This metric effectively captures the peak acoustic energy distribution under pulsed excitation and is well suited for characterizing focused ultrasound fields.

#### Reconstruction of two-dimensional maps from boustrophedon scanning

Two-dimensional planar scans were performed using a row-wise boustrophedon (serpentine) scanning strategy. Because the acquired data form a one-dimensional intensity sequence, the recorded values must be rearranged according to the scanning trajectory to reconstruct the corresponding two-dimensional spatial distribution.

Depending on the scanned plane, the spatial grid is defined as follows:

- For the XY plane, the grid size is *N*_*x*_ × *N*_*y*_, corresponding to the transverse focal cross-section perpendicular to the propagation direction.
- For the XZ plane, the grid size is *N*_*x*_ × *N*_*z*_, corresponding to the axial evolution of the acoustic field along the propagation direction.

The spatial step size was

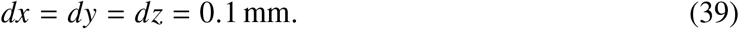

Because serpentine scanning reverses the traversal direction in every even row, horizontal flipping was applied to even-indexed rows during reconstruction to restore the correct spatial ordering. The final two-dimensional acoustic intensity distribution is represented as

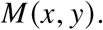

#### Focal spot quantification: two-dimensional FWHM

To quantify the focusing performance, the global maximum position (*x*_max_, *y*_max_) was first identified in the two-dimensional intensity matrix *M* (*x, y*).

Two orthogonal cross-sections passing through the focal peak were then extracted:

- Transverse profile: *M* (*x, y*_max_)
- Longitudinal profile: *M* (*x*_max_, *y*)

The full width at half maximum (FWHM) was computed from the normalized profiles by determining the two positions at which the amplitude decreases to 50% of the peak value. The focal spot size was defined as

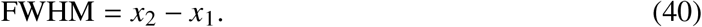

FWHM calculations were implemented using the k-Wave toolbox functions and converted into physical length units (mm). For visualization, the half-maximum line and intersection points were marked on the cross-sectional plots, and the calculated FWHM value was annotated in the figure.

### Spectral analysis and peak pressure evaluation

To analyze the frequency composition and bandwidth characteristics of the acoustic signal at the focus, a fast Fourier transform (FFT) was applied to the filtered time-domain signal *p* (*t*):

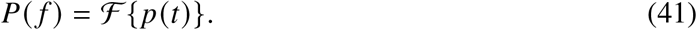

The sampling frequency was *f*_*s*_ = 156.25 MHz.

The single-sided amplitude spectrum was computed as

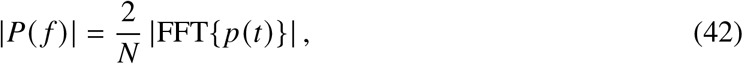

where *N* is the number of sampling points.

To reduce spectral fluctuations, a moving-average smoothing filter was applied to the amplitude spectrum for visualization of the dominant frequency components.

#### Peak pressure evaluation

The peak acoustic pressure was calculated from the unnormalized time-domain signal as

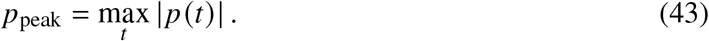

If the oscilloscope voltage signal *V* (*t*) is related to acoustic pressure through hydrophone calibration (e.g., sensitivity *S*), the pressure can be obtained as

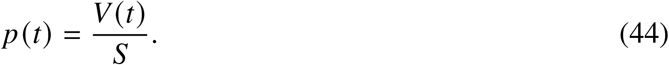

This conversion yields the absolute pressure amplitude at the focus.

### Tests of incident angle, beam offset, and illumination area

To systematically evaluate acoustic field stability and focusing robustness under non-ideal illumination conditions, a custom 3D-printed fiber holder was designed to precisely control laser incidence geometry. The setup allows independent adjustment of three parameters:

#### Incident angle (*θ*)

The angle between the laser propagation direction and the patch normal was adjusted by changing the spatial orientation of the optical fiber. The incident angle *θ* is defined as the angle between the laser beam and the surface normal of the patch. Representative angles (e.g., 0°, 45°, and 75°) were tested to simulate normal and oblique illumination.

#### Lateral beam offset (Δ*x*, Δ*y*)

The fiber position was translated within the patch plane to adjust the lateral displacement of the beam axis relative to the array geometric center. This parameter simulates practical misalignment due to installation error or tissue curvature.

#### Illumination area

The illumination spot size was varied by changing the fiber core diameter or adjusting the distance between the fiber tip and the patch surface, thereby modifying the effective excitation aperture.

These three parameters correspond to illumination direction, position, and coverage area, respectively. By systematically varying their combinations, changes in focal position, focal spot size, and peak pressure were quantified to assess the robustness of the geometrically encoded focusing design against optical misalignment.

### Thermal and long-term stability evaluation

#### Surface temperature monitoring

To assess thermal stability under continuous operation, surface temperature was monitored using an infrared thermal camera (HIKVISION, H21 Pro).

Under continuous laser excitation, surface temperature distributions were recorded over time, and the temperature difference before and after 1 h of operation was compared. Temperature data were extracted from the central region of the patch and from the location of maximum temperature rise to evaluate potential thermal accumulation effects.

This test verifies the thermal safety and material stability of the patch under repeated pulsed excitation conditions.

#### Long-term acoustic stability test

To evaluate the stability of acoustic output under sustained pulsed excitation, long-term operation tests were performed using the following laser parameters:

- Wavelength: 1064 nm
- Single-pulse energy density: 15 mJ/cm^2^
- Pulse duration: 8 ns
- Repetition rate: 10 Hz
- Duration: 1 h

Under these conditions, the photoacoustic patch was continuously excited, and the acoustic pressure signal at the focal position was periodically recorded.

The temporal evolution of the peak focal pressure,

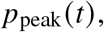

was analyzed to determine whether any amplitude decay or drift occurred during prolonged operation.

Acoustic stability was quantified by comparing the peak pressure at the initial stage of testing with that measured after 1 h of continuous excitation. The relative change in peak pressure was used as a metric to characterize the long-term reliability of the material structure and interfacial bonding.

#### Thermo-acoustic coupling stability evaluation

Thermo-acoustic stability was assessed by jointly analyzing the surface temperature evolution and the temporal variation of focal peak pressure.

If the temperature rise remained minimal and no significant attenuation of focal pressure was observed over time, the device was considered to exhibit stable structural performance and acceptable safety margins within the tested laser energy density range for potential biomedical applications.

## References and Notes

1. A. M. Lozano, et al., Deep brain stimulation: Current challenges and future directions. Neuron 103, 793–806 (2019).

2. J. K. Krauss, et al., Technology of deep brain stimulation: Current status and future directions. Nature Reviews Neurology 17, 75–87 (2021).

3. G. Leinenga, C. Langton, R. Nisbet, J. Götz, Ultrasound treatment of neurological diseases — current and emerging applications. Nature Reviews Neurology 12, 161–174 (2016).

4. J. Blackmore, S. Shrivastava, J. Sallet, C. R. Butler, R. O. Cleveland, Ultrasound neuromodulation: A review of results, mechanisms and safety. Ultrasound in Medicine & Biology 45, 1509–1536 (2019).

5. A. Fomenko, C. Neudorfer, R. F. Dallapiazza, S. K. Kalia, A. M. Lozano, Low-intensity ultrasound neuromodulation: An overview of mechanisms and emerging human applications. Brain Stimulation 11, 1209–1217 (2018).

6. W. J. Tyler, et al., Remote excitation of neuronal circuits using low-intensity focused ultrasound. Nature Neuroscience 14, 168–170 (2011), doi:10.1038/nn.2681.

7. Y. Tufail, et al., Transcranial pulsed ultrasound stimulates intact brain circuits. Neuron 66, 681–694 (2010).

8. W. Legon, et al., Transcranial focused ultrasound modulates the activity of primary somatosensory cortex in humans. Nature Neuroscience 17, 322–329 (2014).

9. H. Estrada, et al., High-resolution fluorescence-guided transcranial ultrasound mapping in the live mouse brain. Science Advances 7, eabi5464 (2021).

10. R. Noureddine, et al., Guidelines for successful motor cortex ultrasonic neurostimulation in mice. Ultrasonics 128, 106888 (2023).

11. C. Pu, et al., A wearable spiral ultrasound patch for focused ultrasound peripheral neuromodulation. bioRxiv (2024).

12. K. W. K. Tang, et al., Bioadhesive hydrogel-coupled and miniaturized ultrasound transducer system for long-term wearable neuromodulation. bioRxiv (2024).

13. V. Pashaei, et al., Flexible body-conformal ultrasound patches for image-guided neuromodulation. IEEE Transactions on Biomedical Circuits and Systems 14, 305–318 (2020).

14. T. Xu, et al., Wearable ultrasound device for long-term neural stimulation. Advanced Science 9, 2201234 (2022).

15. C. Wang, et al., Stretchable ultrasonic transducer arrays for wearable imaging. Advanced Materials 34, 2200159 (2022).

16. W. Du, et al., Conformable ultrasound breast patch for deep tissue scanning and imaging. Science Advances 9, eadh5325 (2023).

17. F. Zou, et al., A wearable spatiotemporal controllable ultrasonic device with amyloid-beta disaggregation for continuous Alzheimer’s disease therapy. Science Advances (2025).

18. L. Shi, et al., A fiber optoacoustic emitter with controlled ultrasound frequency for cell membrane sonoporation at submillimeter spatial resolution. Photoacoustics 20, 100208 (2020).

19. Y. Li, et al., Optically-generated focused ultrasound for noninvasive brain stimulation with ultrahigh precision. Light: Science & Applications 11, 321 (2022).

20. J. Kim, et al., Candle-soot carbon nanoparticles in photoacoustics: Advantages and challenges for laser ultrasound transmitters. IEEE Nanotechnology Magazine 13, 13–28 (2019).

21. P. Beard, Biomedical photoacoustic imaging. Interface Focus 1, 602–631 (2011).

22. M. Xu, L. V. Wang, Photoacoustic imaging in biomedicine. Review of Scientific Instruments 77, 041101 (2006).

23. L. V. Wang, S. Hu, Photoacoustic tomography: In vivo imaging from organelles to organs. Science 335, 1458–1462 (2012).

24. X. Fan, et al., Photoacoustic wavefront engineering. Light: Science & Applications 9, 164 (2020).

25. L. Shi, et al., Non-genetic photoacoustic stimulation of single neurons by a tapered fiber optoacoustic emitter. Photoacoustics (2020).

26. G. Chen, et al., High-precision photoacoustic neural modulation uses a non-thermal mechanism. Advanced Science 11, 2403205 (2024).

27. G. Montaldo, et al., Time reversal acoustics for focusing in complex media. Physical Review Letters 93, 174301 (2004).

28. K. Melde, et al., Holograms for acoustics. Nature 537, 518–522 (2016).

29. M. Fink, Time-reversal acoustics. Physics Today 50, 34–40 (1997).

30. H. Vogel, A better way to construct the sunflower head. Mathematical Biosciences 44, 179–189 (1979).

31. B. E. Treeby, B. T. Cox, k-Wave: MATLAB toolbox for the simulation and reconstruction of photoacoustic wave fields. Journal of Biomedical Optics 15, 021314 (2010).

32. A. E. Siegman, Lasers (University Science Books) (1986).

33. I. T. Jolliffe, Principal Component Analysis (Springer) (2002).

